# Feature selective adaptation of numerosity perception

**DOI:** 10.1101/2024.05.16.594539

**Authors:** Camilla Caponi, Elisa Castaldi, Paolo Antonino Grasso, Roberto Arrighi

## Abstract

Perceptual adaptation has been used to infer the existence of numerosity detectors, which allow humans to quickly estimate the number of objects in a scene. While adaptation was originally thought to affect numerosity perception regardless of the low-level features of the items, a recent study demonstrated that adaptation is more pronounced when the adapting and adapted (test) stimuli share the same color, compared to when they were colored differently. In this study we explored whether such adaptation reduction depends on a novelty effect induced by changes in stimulus features or whether this effect is observed only when implying an identity change of the stimuli. To this aim, we performed six experiments in which numerosity adaptation was investigated in conditions in which adapting and adapted stimuli were either matched or differed for several low-level (color, luminance, shape, and motion) or high-level (letters’ identity, face emotions) features. Numerosity adaptation was consistently observed across all conditions, but it was reduced when adaptor and test differed in color, luminance and shape. However, when stimuli differed in their motion profile, a very salient perceptual change that does not imply a change in items’ identity, adaptation selectivity vanished. Moreover, adaptation selectivity was not observed when items’ identity was changed by spatial rotations of the same stimulus (letters) or when stimuli were matched for the global configuration (face outline) but differed for the arrangement of local features (mouth, nose, eyes). Interestingly, image dissimilarity between test and adaptor, as quantified by Gabor filters simulating a simplified model of the primary visual cortex, nicely predicted the strength of numerosity adaptation across all conditions. Overall, changes in stimulus identity defined by low-level features, rather than novelty in general, determined the strength of the adaptation effects, provided that the changes were readily noticeable. Our findings suggest that numerosity mechanisms may be able to operate on segregated and categorized visual items in addition to the total quantity of the set, with part of the aftereffects induced by numerosity adaptation occurring after feature-binding.

## INTRODUCTION

Humans share with other animal species the ability to make rapid and reasonably accurate estimates of the number of items without relying on serial counting (Dehaene, 2011). This ability is present from a few hours after birth (Anobile, Morrone, et al., 2021; Izard et al., 2009), occurs spontaneously (Caponi et al., 2024; Castaldi et al., 2018, 2020, 2021; Cicchini et al., 2016, 2019) and universally even in individuals with no access to formal schooling (Ferrigno et al., 2017; Piazza et al., 2013). Dedicated numerosity detectors, clearly identified in non-human animals, subserve this fundamental visual property. Electrophysiological studies found that activity of individual cells in the prefrontal and parietal cortices of macaque monkeys and in the endbrain of crows was finely tuned to the numerosities of visual arrays (Nieder, 2016, 2021; Wagener et al., 2018). In humans, with the noticeable exception of a few electrocorticography studies (Kutter et al., 2018; Van Dijk et al., 2022), evidence of the existence of number-selective detectors mainly comes from psychophysical studies using the adaptation technique (Burr et al., 2018). This technique can reveal neural populations that are responsive to the examined feature (e.g., motion direction, orientation, size) and provide insights into the feature’s representational and functional properties. A large body of studies found that numerosity, like many other primary perceptual features, is susceptible to adaptation which elicits aftereffects: exposing individuals to higher numerosities makes the following ones appear less numerous, suggesting the presence of dedicated neural mechanisms for the analysis of numerical quantities (Aagten-Murphy & Burr, 2016; Arrighi et al., 2014; D. Burr & Ross, 2008; Castaldi et al., 2016; Fornaciai et al., 2016; Grasso, Anobile, & Arrighi, 2021; Grasso, Anobile, Caponi, et al., 2021; Grasso et al., 2022; Togoli & Arrighi, 2021; Tsouli et al., 2021). Leveraging on a similar technique, early fMRI habituation studies in humans recorded distance-dependent signal release from adaptation and described tuning curves for numerosity in the intraparietal sulcus (IPS; see Piazza et al., 2004).

Across all these studies, one remarkable characteristic of numerosity adaptation was that it occurred largely independently of changes in low level features and even sensory modality, suggesting that the representation of number may be abstract and a-modal. Indeed, fMRI studies showed that the signal release from habituation in and around the IPS was significantly higher for numerical deviants than for shape deviants (Cantlon et al., 2006) and that it was notation-independent, resulting in similar recovery of the signal irrespective of the numerical deviant being presented in non-symbolic (dot arrays) or symbolic (digits) format (Piazza et al., 2007). Psychophysical studies showed that, in addition to the visual domain, numerosity adaptation also occurs in other sensory modalities, including audition (Arrighi et al., 2014; Grasso et al., 2022; Togoli & Arrighi, 2021) and touch (Togoli et al., 2021). Importantly, several studies have shown that numerosity adaptation generalized across modalities: adapting to auditory sequences influenced the perceived numerosity of sequences of visual stimuli and vice versa (Arrighi et al., 2014), and the same was observed in the tactile modality (Togoli et al., 2021; Togoli & Arrighi, 2021). Furthermore, the adaptation effect is capable of generalizing across motor and sensory domains (Anobile, Arrighi, et al., 2021; Anobile et al., 2016; Togoli et al., 2020) even in the absence of visual experience (Togoli et al., 2020).

While all these studies clearly point to the existence of generalized and a-modal mechanisms tuned to numerosity perception, it might also be ecologically important to have a system capable of segregating different objects within the visual scene that provides independent numerosity estimates for the items of different groups, such as, for example, edible (ripe) and not edible (unripe) fruits. Indeed, previous evidence showed that human adults can attentively encode, in parallel, different subsets of items defined by color, suggesting that approximate numbers can be stored as a feature of each set (Halberda et al., 2006). In line with the possibility of creating parallel representation of numerosity, a recent study demonstrated that numerosity adaptation is tuned to specific features of the stimuli such as color in vision and pitch in sound (Grasso et al., 2022). Participants were adapted to high numerosity dot arrays with items’ color either matched to or different from the test. Numerosity underestimation (about 25%) was observed for matched colors (either physical or perceived), while the adaptation effect was much reduced when the adaptor and test colors differed, with a similar selective adaptation effect also occurring in audition.

To investigate further the phenomenon of the selectivity of numerosity adaptation, in the current study we assessed which changes in stimulus features lead to adaptation reduction and whether such adaptation reduction is driven by an automatic regain of attention to a novel stimulus (changes in stimulus novelty) or whether it depends on the visual system categorizing stimuli on the basis of their “identity”. In the latter case, numerosity adaptation would be “selective” in that the prolonged exposure to a given class of stimuli would distort the perceived numerosity of items belonging to a different class less. To test these hypotheses, we conducted six experiments wherein adult participants were adapted to high numerosities and tested with stimuli that either matched or differed from the adaptor in various low-level features (color, luminance, shape and motion) or high-level characteristics (letters’ identity, face expressions). The first three experiments modulating color, luminance and shape congruency between adaptor and test stimuli would allow us to replicate previous findings by Grasso et al. (2022) and extend these results to other perceptual domains. In the fourth experiment, we modulated items’ motion profiles (still vs randomly moving items) between adaptor and test, a feature change that provides a striking difference between the stimuli though no change to the items’ identity. In other words, if numerosity adaptation selectivity depends on stimulus identity, and not on its novelty, distortions of perceived numerosity should also occur when observers are adapted to the numerosity of moving items and are then required to estimate that of completely still displays. Next, we investigated whether the numerosity adaptation selectivity can also be observed when stimulus identity is defined by high-level semantic meaning. To this aim we used letters as these allowed us to disentangle stimulus low-level features (shape) from their identity (defined at the semantic level), with different letters obtained by a simple spatial rotation of the same shape (i.e., ‘b’ vs ‘q’). Letters were also useful to test the opposite condition in which the same item identity was preserved despite salient changes in the stimuli low level characteristics such as when the same letter is displayed in different notations (i.e., lowercase ‘b’ vs uppercase ‘B’). If adaptation selectivity is determined by high-level semantic stimulus identity, we should observe the same adaptation effect for ‘b’ and ‘B’ and an adaptation reduction for all the other, different, letters. On the contrary, if adaptation selectivity does not apply to high-level semantic identity, the change in letter notation from ‘b’ to ‘B’ should elicit the weaker adaptation effect, because this pair exhibits the greatest variation in low-level features (such as shape). Finally, in a sixth experiment, the role of stimuli identity was tested via salient ecological stimuli such as faces (represented by smiley ideograms) with different emotional expressions (happy or sad faces) or with corrupted spatial arrangement of the inner components (scrambled faces). If adaptation selectivity is related to stimulus identity, we should observe the same adaptation effect irrespective of faces’ emotion, unless these are coded as different individuals (e.g. enemies vs friends). On the contrary, we expect a decreased adaptation for scrambled compared to non-scrambled faces since they should be perceived as belonging to different classes of stimuli. To anticipate the results, we found that stimulus identity, as defined by low-level features, is the crucial factor in determining the magnitude of numerosity adaptation aftereffects, while complex high-level characteristics pointing to semantic or emotional cues were found to be much less effective in modulating the magnitude of adaptation aftereffects. Using a model of the primary visual cortex response to different classes of stimuli used across our experiments, we measured the low-level “dissimilarity” between adaptor and test stimuli and found that such parameters strongly affected the adaptation magnitude, with higher dissimilarity being associated with weaker adaptation aftereffects.

## METHODS

### Participants

The sample size was calculated using an a priori Power analysis (G*Power software, version 3.1, Faul et al., 2007). The effect size was estimated from the study of Grasso et al., (2022) that exploited very similar methods. With an α = 0.05 and a Power of 0.9, the analysis suggested a required sample size of 18 participants.

Across all experiments, a total of 56 participants took part in the study (mean age: 25.2, standard deviation: 3.9, 37 females). Each participant performed one or more of six different experiments which differed in the type of feature characterizing the arrays of visual stimuli as detailed below. All participants had normal or corrected-to-normal visual acuity and provided written informed consent prior to the experiment. The research was approved by the ethics committee (Commissione per l’Etica della Ricerca, University of Florence, July 7, 2020, n. 111) and it was performed in accordance with the Declaration of Helsinki.

### Stimuli and Procedure

Participants sat 57 cm away from a 27″ screen monitor (resolution 2560×1440 pixel; refresh rate 60 Hz), in a quiet and dimly lit room.

Across six experiments we tested the selectivity of numerosity adaptation to different classes of stimuli where adapting and adapted items were congruent or not congruent for color (Exp 1), luminance (Exp 2), shape (Exp 3) and motion (Exp 4). Two additional experiments were carried out to test selective adaptation for conditions in which stimuli identity prompted a semantic recognition process such as letter identification (Exp 5) and whether adaptation selectivity occurred for high level stimuli (stylized faces) that differed in emotional state or arrangement of the local inner components (faces vs scrambled faces, Exp 6).

For each experiment, we used a two-alternative forced-choice method (2AFC) to estimate participants’ perceived numerosity, separately in baseline and adaptation conditions (see Figure 1). Stimuli were arrays of non-overlapping items drawn within a virtual circle (diameter: 8 deg), displayed briefly (200 or 500 ms, depending on the experiment) and simultaneously to the left or to the right of a central fixation point (eccentricity: 10 deg), on a mid-grey background. One of the two arrays was the reference with numerosity fixed at 24 items, while the other stimulus varied in numerosity for each trial between 10 and 60 or 12 and 48 depending on the experiment. In the adaptation condition, the reference and test stimuli were preceded by an adaptor which was presented for a longer period (2 or 4 sec, depending on the experiment) in the same position as the test and with a fixed higher numerosity (48 or 72 items, depending on the experiment). In the experiments with colored and achromatic dots and letters, the test and reference stimuli (12-48 items) were presented for 200 ms, while the adaptor stimulus, consisting of 48 items, lasted 2 seconds. In the experiments with moving dots, faces, and shapes, the test and reference stimuli (12-48 items for moving dots and 10-60 items for faces and shapes) were presented for 500 ms and the adaptor, consisting of 72 items, lasted 4 seconds. In each trial, regardless of the kind of stimuli exploited, the task for the participants was to indicate, via key press, whether the stimulus on the left or right was the more numerous.

**Figure 1.**
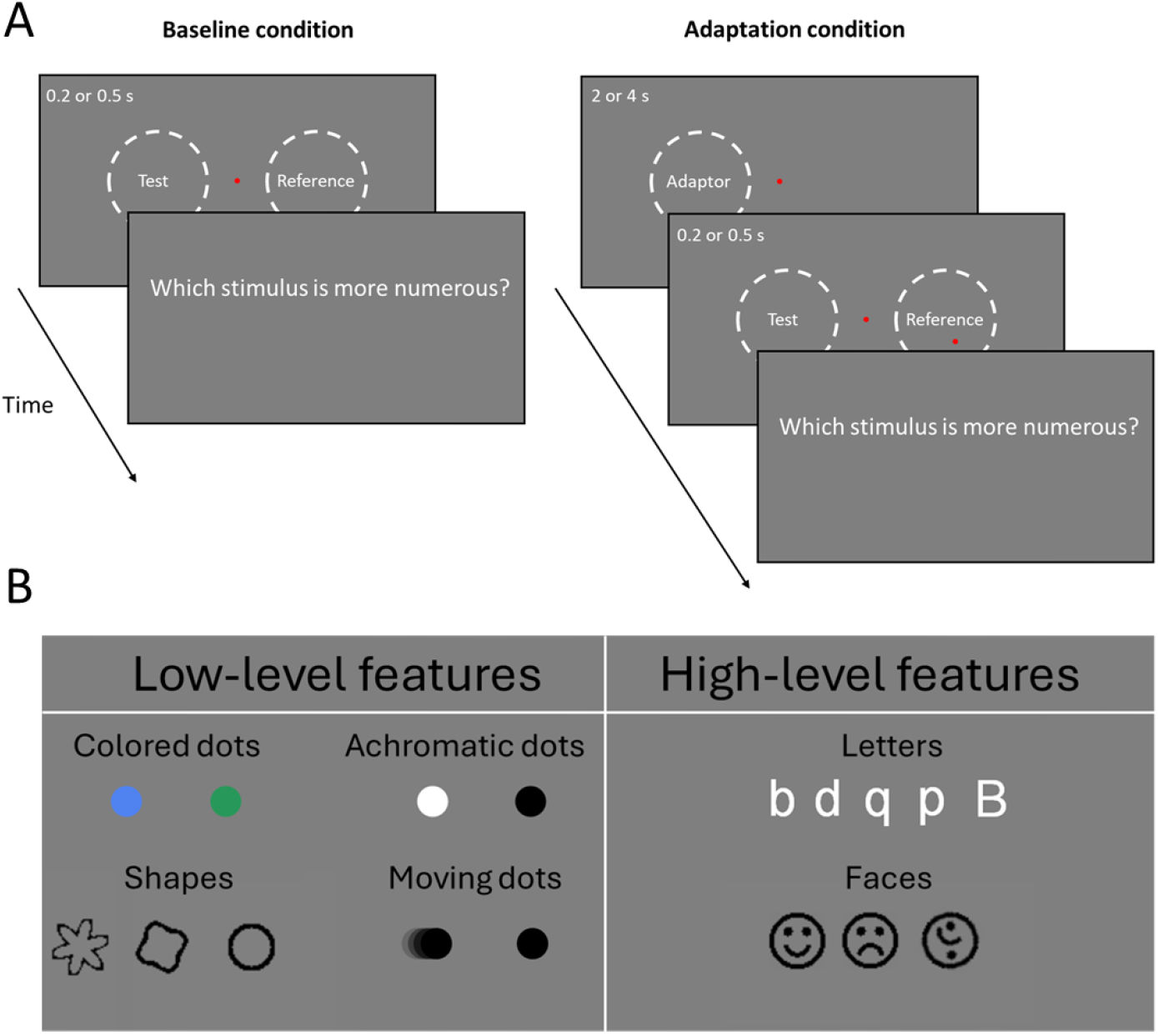
Overview of the experimental design and stimuli. (A) The panels represent the time course of the two conditions, baseline and adaptation. (B) Examples of the stimuli used across the six experiments.

Each experiment aimed at investigating whether and to what extent the congruency between the adapting and adapted stimuli for a given feature modulated the magnitude of the adaptation aftereffects. To this end, each session comprised trials in which adaptor and test were identical (congruent condition) and trials in which they differed for a given feature (incongruent condition) with the two conditions presented interleaved. The difference in adaptation magnitude measured in the congruent and incongruent conditions defined the amount of selectivity of numerosity adaptation. Below is a detailed description of the stimuli used in each of the six experiments.

#### Experiment 1 - Color. 20 participants took part in this experiment

Stimuli were arrays of colored dots (diameter: 0.4 deg). In both the baseline and adaptation conditions, the color of the test and reference stimuli randomly varied between blue (RGB code: 80 130 240) and green (RGB code: 40 152 90), while the color of the adaptor was kept fixed (blue).

#### Experiment 2 - Luminance. 20 participants took part in this experiment

Stimuli were arrays of black and white dots (diameter: 0.4 deg). The color of test and reference randomly varied between black and white, while the color of the adaptor was fixed to white. In the adaptation condition, in order to avoid visual aftereffects elicited by the prolonged presentation of the white adaptor, a full screen mask composed of randomly arranged black and white squares (0.2x0.2deg), was briefly (20ms) presented soon after the adaptor and before the presentation of the test and reference.

#### Experiment 3 - Shape. 20 participants took part in this experiment

Stimuli comprised arrays of three different shapes. One of the shapes was a circle with a diameter of 0.8° while the other two shapes were obtained by sinusoidally modulating the radius of the circle over 360°: when the modulation was 0.5 deg in depth and repeated 4 times, the patches resembled a distorted square while with a 0.8 deg in depth modulation repeated 6 times we obtained a stimulus resembling a flower. Importantly, because the positive and negative modulation of the radius were defined by a symmetric sinusoidal profile, these were identical so the perimeter and the area of all the patches were perfectly matched despite their different appearances. In each trial, regardless of the shape of the stimuli, half of them were colored in white and half in black to match the average luminance stimuli with that of the mid gray background.

#### Experiment 4 - Motion. 20 participants took part in this experiment

Stimuli comprised arrays of moving and static dots (diameter: 0.8 deg), half white and half black. In the baseline condition, dots in the test and reference stimuli were either static or moving, while the adaptor always consisted of an array of moving dots. All dots moved within an invisible area of 8° diameter at a constant speed of 7 deg/sec across a gray background. They moved in straight lines and, when colliding with other dots or the boundaries of the conscription area, they bounced appropriately (obeying the laws of physics).

#### Experiment 5 - Letters. 26 participants took part in this experiment

Stimuli comprised arrays of different letters. For the test and the reference stimuli we used letters ‘b’, ‘p’, ‘d’, ‘q’ and ‘B’, while the adaptor only included ‘b’s. All letters were matched in size and subtended around 1.2 deg.

#### Experiment 6 - Faces. 20 participants took part in this experiment

Stimuli comprised arrays of stylized faces (smileys). Test and reference randomly varied between smiling faces, sad faces and scrambled faces, while the adaptor always consisted of an array of smiling faces. Each stimulus subtended 0.8° by 0.8° and, in each display, half of the faces were white and half black to match as average luminance the mid gray background.

In the experiments including shapes, letters and faces, we added a secondary task to ensure participants were actually capable of discriminating the different stimuli from each other: in one third of the trials (chosen at random), after the response to indicate the most numerous stimulus, participants were required to also indicate - via key press-what class of stimuli (in terms of color, shape, letter) they have been presented with in that trial.

Stimuli were generated and presented with PsychToolbox 3.0.16 routines (Brainard, 1997; Kleiner, 2007) for Matlab (Ver. R2021b, The Mathworks, Inc http://mathworks.com)

### Data Analysis

Data were analyzed separately for each participant and experiment. For each condition, we plotted the percentage of trials in which the test stimulus was perceived more numerous than the reference as a function of tested numerosities and fitted data distribution with a best-fitting cumulative gaussian function. The 50% point of the function defines the point of subjective equality (PSE), the physical numerosity at which the test stimulus was perceptually matched to the reference. We calculated the adaptation effect by subtracting, for each participant, the PSE for the adaptation condition from the PSE measured in baseline (no adaptation) and normalizing this value by the latter.

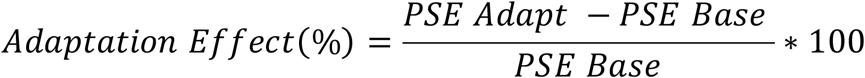

For experiments involving shapes, letters, and faces, we also calculated the percentage of correct responses in the secondary task wherein participants indicated the stimulus identity.

Data were analyzed with paired sample t-tests, Repeated-Measures ANOVAs and Bonferroni corrected post-hoc t-tests (with p-values corrected for multiple comparisons reported as p_bonf_). Bayes factors are expressed as Log10 Bayes factors (LogBF10). Positive values of LogBF10 are conventionally interpreted as providing substantial (0.5–1), strong (1–2) or decisive (> 2) evidence in favor of the alternative hypothesis. Effect sizes (η^2^ or Cohen’s d) were also reported. Statistical analyses were performed using JASP (version 0.16.1, The JASP Team 2022, https://jasp-stats.org/).

### Image dissimilarity analysis

To achieve an objective parameter with which to assess the similarity amongst the stimuli used in the six experiments, we calculated the image dissimilarity between all items exploited as adaptors and tests. To this aim we used the image dissimilarity toolbox developed by Seibert and Leeds using the Gabor Filterbank method (Leeds et al., 2013). This method projects each image onto a Gabor wavelet pyramid simulating a simplified model for primary visual cortex (see Kay et al., 2008). The filters span eight orientations and four sizes across the image. The Euclidean distance is used to calculate the difference between the vectors of filter responses associated with each pair of images. Leeds and coll. found that the representational dissimilarity matrix resulting from the Gabor filter bank model applied to a dataset of naturalistic images positively correlated with neural activity of the occipital areas, but also with that of ventrotemporal and dorsal parietal cortices (Leeds et al., 2013). The distance matrix correlations resulting from the Gabor filter bank model were correlated with the adaptation difference between the congruent condition and the incongruent conditions, across all experiments. The correlation was reported using the Spearman correlation coefficient (rho).

## RESULTS

We first investigated the selectivity for numerosity adaptation due to color with the aim of replicating the finding by Grasso et al. (2022). Participants were adapted to arrays of blue dots and then discriminated which of two simultaneously presented arrays displayed, one on the left and one on the right, was more numerous. When the dots of test and reference stimuli were blue, like the adapter, the condition was congruent, when they were displayed in green the condition was incongruent. Figure 2A shows the psychometric functions averaged across participants for numerosity discrimination separately for baseline (no adaptation) and adaptation conditions. As the PSE measured in the two baseline conditions, one comprising only blue and the other only green dots, was not significantly different (t(19)=-0.49, p=0.63), we pooled these data together to yield a single baseline condition (black curve). The red curve shows the condition where test and reference matched the adaptor for color (congruent condition) while the blue curve shows the condition where test and reference had a different color from the adaptor (incongruent condition). The larger rightward shift of the red psychometric curve indicates a larger adaptation effect when the adapting and adapted stimuli did look identical (congruent compared).

**Figure 2.**
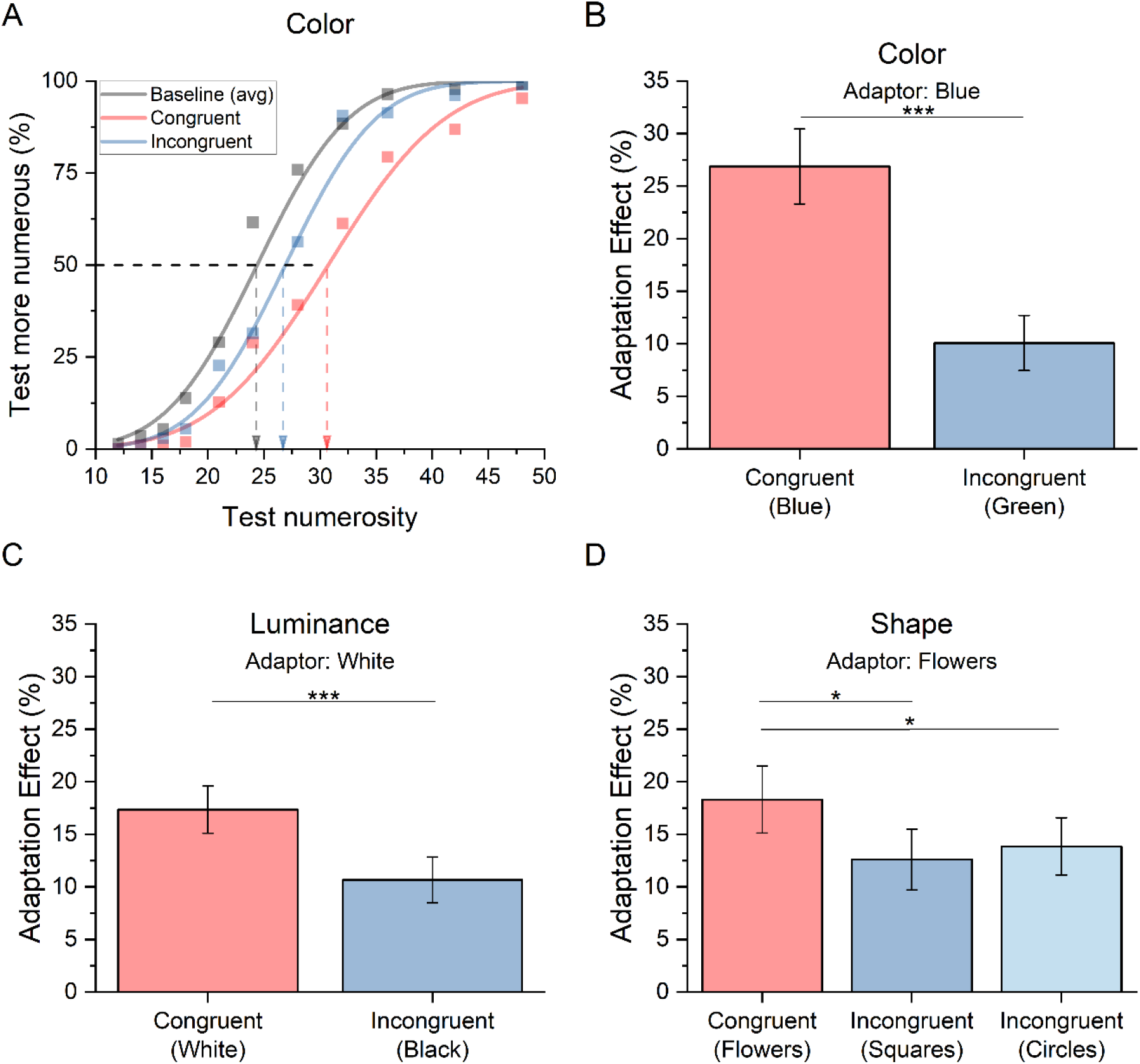
Numerosity adaptation effect measured in the first three experiments in which congruency between adaptor and test was manipulated for color, luminance or shape. (A) The functions plot the percentage of trials where the test appeared as more numerous than the reference as a function of test numerosity (shown in the abscissa). The black curve shows the aggregate psychometric function measured in baseline while the red and blue curves indicate adaptation for the congruent and incongruent conditions respectively. The vertical dashed arrows show the estimates of the PSE in each condition. The bar graphs on the right panel show the mean adaptation effect (%) measured in the congruent (red) and the incongruent (blue) conditions when the manipulated feature was dots’ color (B). Magnitude of adaptation effects for congruency between adaptor and test for luminance (C) and item shape (D). Error bars correspond to 1 s.e.m.

A paired sample t-test confirmed that the percentage of adaptation effect was significantly larger in the congruent compared to the incongruent condition (t(19) = 5.61, p <0.001; Cohen’s *d* = 1.3; LogBF10 = 3.1, Figure 2B), replicating previous results by Grasso et al (2022).

Next, we investigated whether numerosity adaptation selectivity extends to perceptual features other than color, and measured the adaptation magnitude for clouds of achromatic dots that could be either white or black. The result (Figure 2C) showed a similar trend to that obtained for the manipulation of color, that is, a greater adaptation effect in the congruent condition (adapting and adapted stimuli were all white) compared to the incongruent condition (white adaptor and black test and reference). A paired sample t-test confirmed that the difference between the adaptation effects was significant (t(19) = 3.9, p =0.001; Cohen’s *d* = 0.9; LogBF10 = 1.6) extending the selectivity of numerosity adaptation to luminance congruency.

In a third experiment, we aimed to establish whether the selectivity of numerosity adaptation, previously found for differences in color and luminance, extended also to another feature typically used to determine object’s identity: its shape. In the congruent condition the shape of the test and reference stimuli matched that of the adaptor (‘flower shape’), while in the incongruent conditions they looked completely different, resembling either a square or a circle. Figure 2D shows that, again, the adaptation effect was stronger for the congruent compared to the incongruent condition. A Repeated-Measure ANOVA revealed a significant effect of item shape on adaptation (F(2,38)=4.3, p=0.02, η^2^=0.19). Post-hoc t-tests revealed that the two incongruent conditions both differed from the congruent condition (squares: t(19) =2.6, p_bonf_=0.043, Cohen’s *d*= 0.50; circles: t(19) =2.5, p_bonf_=0.047, Cohen’s *d*= 0.50), but not from each other (t(19) =-0.04, p_bonf_=1, Cohen’s *d*= - 0.01).

Interestingly, while color, luminance and shape can define objects’ identity, their change between adapting and adapted stimuli also introduced a “novelty effect” that might be able to disrupt part of the adaptation effect. To assess whether stimuli identity or novelty determined the reduction in adaptation effect in the incongruent condition, we modulated the congruency between adapting and test/reference stimuli in terms of their motion (moving vs still stimuli). If numerosity adaptation selectivity depends on stimulus identity, we expected the same adaptation effect regardless of the motion congruency as it is more common for an object to change position over time than change its color or shape. Figure 3A shows the psychometric functions, averaged across participants, for the conditions in which the adapting displays consisted of clouds of moving dots and in which dots of the test and reference were moving or not (red and blue respectively). The black curve represents the baseline condition with no adaptation. In the baseline condition, data for static or moving dots were pooled, since there was no difference in numerosity perception (PSE, t(19)=0.7, p=0.5). The rightward shift of the blue and red psychometric curves indicates a robust numerosity adaptation effect. However, as it is clear from inspection, both curves are rather superimposed, suggesting that adaptation magnitude was not affected by motion congruency between test and adaptor. A paired sample t-test confirmed that there was no significant difference between adaptation in the congruent and incongruent conditions (t(19) = 0.50, p = 0.6; Cohen’s *d* = 0.11; LogBF10 = -0.60), indicating that adapting to moving stimuli distorts the perceived numerosity of both, moving and still stimuli to the same extent (Figure 3B).

**Figure 3.**
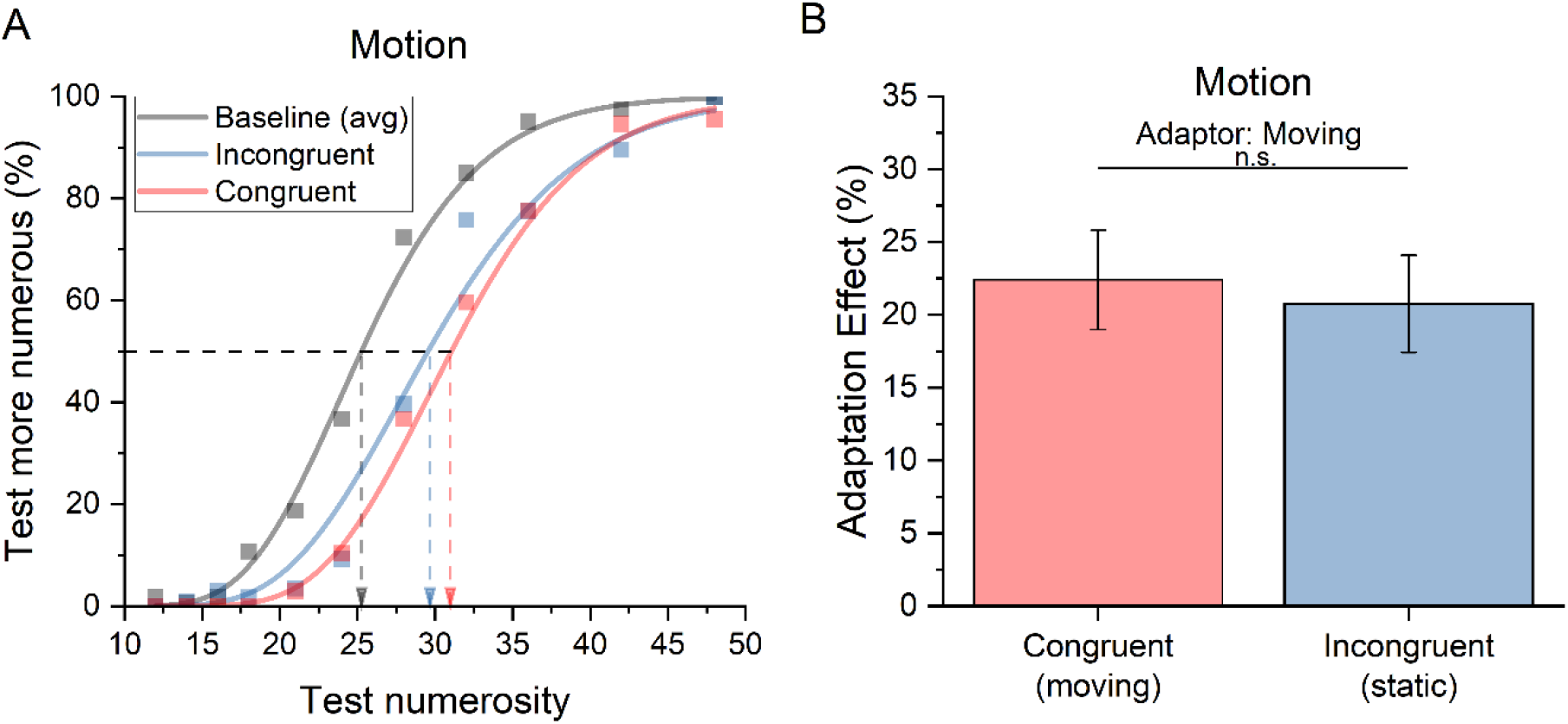
Numerosity adaptation effect measured in the fourth experiment in which the motion of adapting and adapted dots was manipulated. (A) The functions plot the percentage of trials where the test appeared more numerous than the reference as a function of the test numerosity (shown in the abscissa). The black curve shows the aggregate psychometric function measured in the baseline (no adaptation) while the red and blue curves indicate the congruent and incongruent conditions respectively. The vertical dashed arrows show the estimates of the PSE in each condition. Bar graphs show the mean adaptation effect (%) measured in the congruent (in red) and the incongruent (in blue) conditions. Error bars represent 1 s.e.m.

Given that all previous experiments seem to suggest that the congruency of stimuli identity between adapting and adapted stimuli might play a key role in defining the magnitude of numerosity adaptation, we designed two additional experiments to test whether the same applied when items’ identification prompts a high-level, semantic process. To this aim we used letters as they allow us, by means of simple spatial rotations of the very same shape, to create different letters (i.e. from p to b). In other words, by using letters, it is possible to create conditions wherein minimal changes in the low-level characteristics of stimuli result in entirely different high-level semantic meanings. Moreover, as the same letter can be written in different formats, either capital or lower-case, it is possible to achieve bold changes in the shape of the stimuli while maintaining the same high-level semantic meaning (or identity). We thus defined a congruent condition in which the test and reference were identical to the adaptor (‘b’). Then we designed three incongruent conditions in which the high-level semantic change was maximized (by the letter change) while keeping minimal the alteration of stimuli low-level features (‘d’, ‘p’, ‘q’). Finally, we devised an additional incongruent condition in which changes in the low-level features were maximized, but the letter identity was maintained congruent between test and reference (‘B’) and the adaptor (‘b’). As shown in Figure 4A, a strong and robust adaptation effect was observed across all conditions, irrespective of what kind of letter was displayed in lower-case.

**Figure 4.**
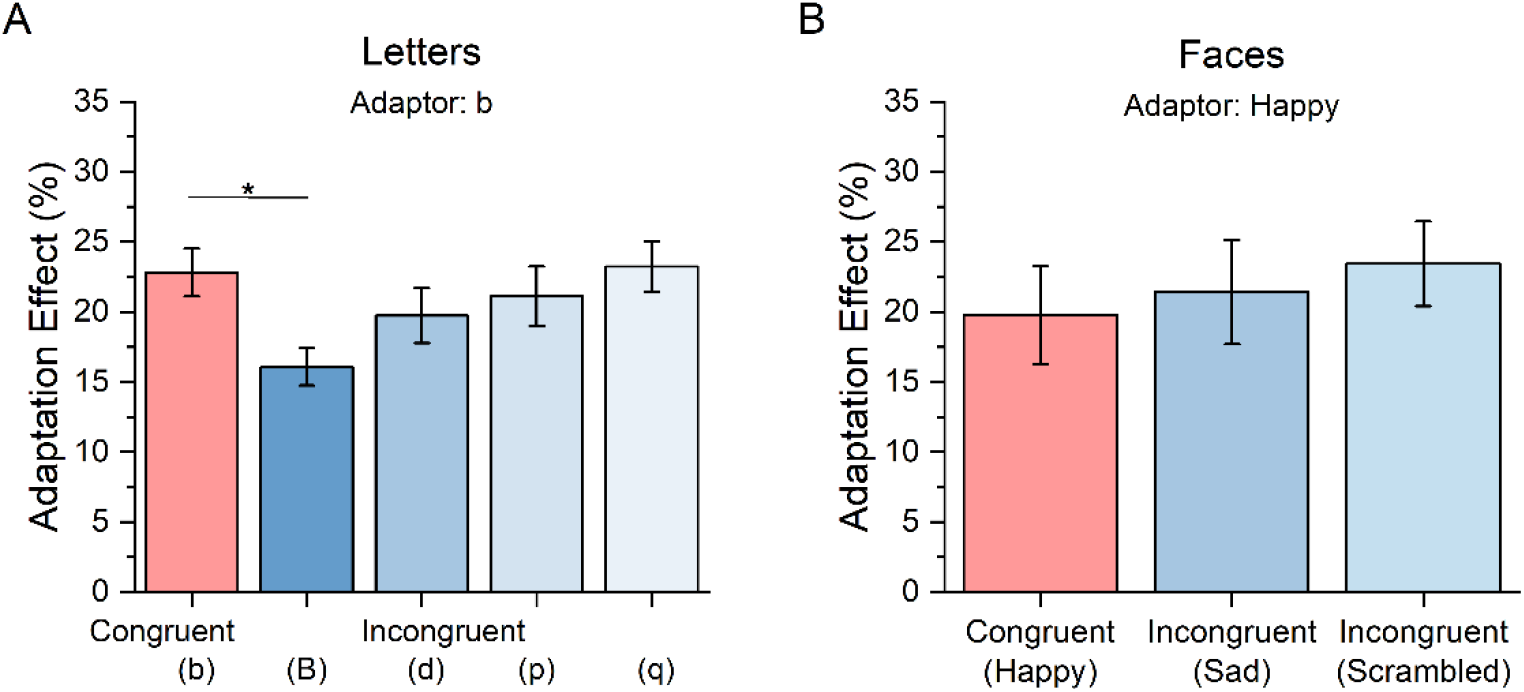
Numerosity adaptation effect measured in the fifth and sixth experiments in which congruency between adaptor and test was manipulated with respect to high level features. Bar graphs show the mean adaptation effect (%) for the congruent (in red) and the incongruent (in blue) conditions, separately for (A) letters and (B) faces. Error bars represent 1 s.e.m.

However, adaptation was remarkably decreased when the test stimulus contained ‘B’s. A Repeated-Measure ANOVA revealed an effect of the letter on adaptation (F(4,25)=4.9, p=0.001, η^2^=0.17). Specifically, post hoc t-tests indicated that only when the incongruent condition consisted in a font/shape change (from ‘b’ to ‘B’), the adaptation effect was lower than the congruent condition (t(25) =3.7, p_bonf_=0.004, Cohen’s *d*= 0.75), while in all other conditions, the adaptation magnitude was the same (all p-values not significant). This result suggests that adaptation selectivity is less determined by the high-level semantic meaning (i.e. letters identity) than by the low-level changes of the stimuli features such as the changes in shape induced by changing the letters’ font.

Finally, to manipulate stimuli similarity via ecological stimuli containing both global and local information, in a sixth experiment we adapted and tested participants with stylized representations of faces. In the congruent condition, test and reference were identical to the adaptor (smiling faces or ‘smileys’), while in the incongruent condition test and reference stimuli consisted either of ‘sad faces’, achieved by rotating upside down the mouth, or ‘scrambled faces’ in which the internal elements (mouth, nose and eyes) were rearranged from their canonical positions. Figure 4C shows that in this experiment, we achieved consistent adaptation aftereffects in all conditions regardless of stimuli similarity (repeated-Measure ANOVA; F(2,38)=0.55, p=0.58, η^2^=0.03). Therefore, when stimuli similarity was just defined by changes of local elements while the same global configuration remained the same (faces outline), no categorization of the stimuli seems to take place to yield selective effects of numerosity adaptation.

Figure 5A summarizes the results of all six experiments. It is clear from inspection of all 6 experiments that we found a significant adaptation effect also for the incongruent conditions, suggesting a consistent part of the numerosity adaptation aftereffects occurs independently of the characteristics of the displayed items. This suggests that stimuli categorization by similarity might modulate but not completely account for numerosity adaptation. However, Fig 5A also clearly shows that when changes between adaptors and test stimuli regarded low-level features (e.g. color, luminance, shape or font) a significant reduction of the adaptation effects in the incongruent conditions compared to the congruent condition is observed. Intriguingly, such a selective effect did not hold true in the case of motion as adapting to moving stimuli also robustly distorted numerosity perception of static stimuli. Similarly, the congruency between adaptor and adapted stimuli did not play a significant role when stimuli were categorized based on complex, high-level semantic features (e.g. faces or letters). Is it possible that the conditions in which we failed to find evidence for a selectivity in numerosity adaptation were those in which the observers were not able to distinguish between the stimuli similarity of adaptor and test? Our data suggest that this explanation is quite unlikely. Indeed, as reported in the methods section, to specifically control for items detectability we added an additional question for the participants in three experiments (shapes, faces and letters) where the stimuli were more complex. In one third of the trials participants were required to indicate what kind of item was displayed after having indicated the more numerous stimulus. Accuracy in this secondary task was found to be almost perfect in all experiments, with the percentage of correct response around 90-95% in all conditions.

**Figure 5.**
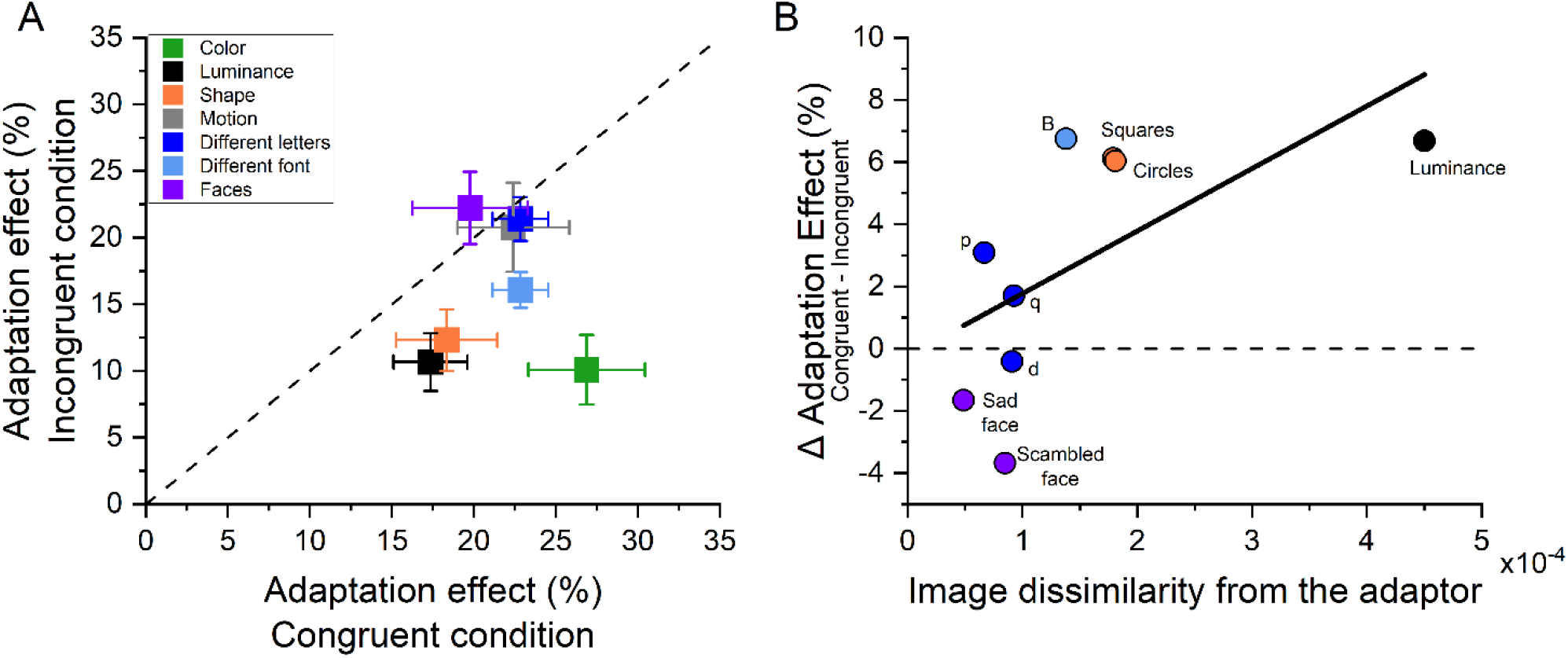
Summary of the adaptation effects measured across the six experiments. (A) Scatter plot of adaptation effects in the congruent (on the x-axis) and incongruent (on the y-axis) conditions, across experiments. Datapoints below the bisector, indicate a stronger adaptation effect in the congruent compared to the incongruent condition. (B) Correlation between the image dissimilarity (on the x-axis) and the adaptation effect difference between the congruent and incongruent conditions (on the y-axis). The dashed horizontal line indicates the same adaptation effect between the congruent and incongruent conditions.

Even though the observers were capable of identifying the elements included in the test and reference and thus to detect whether these were congruent or incongruent relative to the items displayed in the adaptor, there is still the possibility that some feature changes were just more salient than others, with the most striking features changes prompting a more efficient items categorization that, in turn, might have affected numerosity adaptation selectivity. To test this possibility we calculated the image dissimilarity between adaptor and test stimuli by using a simplified model for the primary visual cortex (see Methods for details). Then, we correlated the representational dissimilarity matrix resulting from a Gabor filter bank model with the difference of the adaptation magnitude between the congruent and the incongruent conditions.

Figure 5B shows that as image dissimilarity between the adaptor and test stimuli increased so did the difference between congruent and incongruent adaptation effects (ρ = 0.75 p = 0.025), pointing at image dissimilarity as a possible key factor in determining the strength of the adaptation effect.

## DISCUSSION

In the current study, we found that numerosity adaptation is a flexible mechanism. Despite the fact that adaptation aftereffects have been observed across a wide variety of different conditions, and regardless of the kind of stimuli used as adaptor or test, our experiments clearly demonstrate that, under some circumstances, numerosity adaptation is significantly larger when the adapting and test stimuli are matched for non-numerical low-level features. For example, numerosity underestimation following adaptation to high numerosity was larger when the test matched the adaptor for color, luminance, or shape compared to when the stimuli differed. Stimulus novelty alone was not sufficient to explain the selectivity of numerosity adaptation, as adaptation to moving stimuli also affected the perceived numerosity of still stimuli, despite the change in the motion profile being very salient. This result seems to suggest that adaptation selectivity might arise from a classification process capable of categorizing the adaptor and test stimuli, with the largest adaptation aftereffects occurring when stimuli shared the same features. Interestingly, this would set the similarity between adaptation and another contextual effect, serial dependency, in which the perception of a stimulus in any given moment is biased to look more similar to that presented a little earlier. Indeed, it has been reported (Cicchini et al., 2017; Fischer & Whitney, 2014) that the more similar two stimuli are to each other, the more the attractive effect between the stimuli is observed. Similarly, we demonstrated here that the more similar adapting and adapted stimuli look to each other, the more distorted the numerosity estimates of the latter were. Despite it not being clear which conditions yield attractive or repulsive effects, finding a similarity between the process of adaptation and serial dependency might help to shed light on the perceptual mechanisms underpinning these effects. However, not all the differences between adapting and adapted stimuli were able to attenuate adaptation effects. In numerosity tasks in which letters were used as stimuli, simple spatial rotations of the same visual pattern yielding different semantic meanings (different letters) failed to induce adaptation selectivity. On the contrary, keeping the letter the same but changing the format (from lower to uppercase) provided a clear reduction of adaptation aftereffects, indicating that differences on low level features of the stimuli are capable of overruling congruency at the semantic level. Moreover, when stimuli were matched for global information (outlines of stylized faces) but differed for local information (position of the faces components) no adaptation selectivity was found, indicating that the stimuli categorization process bound to numerosity mechanisms might just be able to make a coarse analysis of items’ features. Taken together, our results reveal that the magnitude of adaptation aftereffects is dependent on the low-level image dissimilarity between adaptor and test with such dissimilarity being quantitatively estimated by a simple model simulating V1 information processing.

It is interesting to note that our results suggest that numerosity perception operates in parallel for different groups of objects and that the perceived numerosity of each group can be independently adapted to some extent. From an ecological perspective it would be advantageous to form different and selective representations of the number of target objects in a visual scene. Importantly, not all low-level salient changes might be relevant: an object may change its position, but it rarely changes its physical properties such as color or shape. The adaptation selectivity observed for color, luminance, and shape, but not motion, suggests that numerosity perception is sensitive to salient environmental low-level features which allow us to segregate objects into categories. Numerosity mechanisms might be sensitive to constancies in these features which can be susceptible to selective adaptation. This is in line with previous evidence showing that we can simultaneously keep track of numerosity up to three different subsets of items defined by color (Halberda et al., 2006). Further evidence, suggesting that numerosity mechanisms operate on segregated and categorized visual items, comes from studies showing that various grouping cues can alter numerosity perception. The precision of our numerical estimates increases when multiple items can be segregated in small groups, a phenomenon called “groupitizing” (Anobile et al., 2020; Caponi et al., 2023; Ciccione & Dehaene, 2020; Maldonado Moscoso et al., 2020; Starkey & McCandliss, 2014). When items are grouped by connectedness or symmetric cues, a strong underestimation of perceived numerosity is achieved, suggesting that numerosity mechanisms spontaneously inform us of the number of groups of objects (Anobile et al., 2017, p. 20; Apthorp & Bell, 2015; Castaldi et al., 2021; Fornaciai et al., 2016; Franconeri et al., 2009; He et al., 2009; Maldonado Moscoso, Anobile, et al., 2022). Previous studies showed that under challenging dual tasks, the influence of grouping cues on numerosity perception is much reduced. Depriving visual attention strongly decreased estimation precision of grouped arrays (Maldonado Moscoso et al., 2020), as well as both the symmetry- and the connectedness-induced numerosity underestimation (Maldonado Moscoso et al., 2023; Pomè et al., 2021), suggesting that the grouping process requires attention. Neuroimaging studies further corroborate this, revealing that grouping cues modulate EEG components typically associated with visuo-spatial attention (Caponi et al., 2023; see reviews by Hillyard & Anllo-Vento, 1998; Mangun & Hillyard, 1991) and the pattern of activity of areas starting from V3 and beyond (Fornaciai & Park, 2018; Maldonado Moscoso, Greenlee, et al., 2022). By showing that numerosity adaptation selectively impacts different categories of items, the current study suggests that it might occur after grouping or feature bounding, presumably at a relatively late attention-dependent stage. This is in line with previous studies showing that numerosity adaptation operates on perceived rather than physical numerosity when the two differ due to perceptual grouping (Fornaciai et al., 2016) and that the availability of visuospatial attention during the adaptation period crucially determines the adaptation aftereffects (Grasso, Anobile, & Arrighi, 2021; Grasso, Anobile, Caponi, et al., 2021).

Clearly, grouping cues have to be sufficiently salient to be perceptually noticeable. In a previous paper Burr and Ross (2008) did not find adaptation selectivity to orientation, and size changes between adaptor and test. There are at least two explanations for this observation. First, size and orientation changes may not necessarily correspond to changes in objects identity: the same object appears larger when it approaches the observer and can change orientation. An alternative explanation for this discrepancy is that in the experiment by Burr and Ross (2008), the changes were not sufficiently salient to be easily discriminated in the peripheral vision. In the current study we demonstrated that detectability of stimulus changes is indeed a key factor in determining the strength of the adaptation effect, by showing that it can be predicted by image dissimilarity. Future studies should better investigate how size and orientation changes impact the adaptation selectivity.

The reduced discriminability of stimulus changes might in part explain why adaptation selectivity did not occur when adaptor and test differed for high-level features. Adapting to the letter ‘b’ generalized to all the other lower-case letters and adapting to stylized happy faces generalized to both sad and scrambled faces. The dissimilarity between the adaptor and test stimuli in these cases was very low and likely explains the generalization of adaptation. However, the fact that adapting to the letter ‘b’ did not generalize to the capital ‘B’ suggests that the low-level shape change, prevailed on letter identity in determining the adaptation selectivity or that high-level semantic grouping, might not influence numerosity perception, at least for non-ecological culturally mediated stimuli, such as letters. For now, we can conclude that, at least with the stimuli currently used, the low-level feature changes prevailed on the high-level feature changes in driving the adaptation selectivity.

The fact that the selectivity of the adaptation effect can be predicted by image dissimilarity estimated by a simple model of V1 does not necessarily mean that adaptation occurs at this level. Indeed, image dissimilarity calculated with this model also correlated with activity in many areas along the ventral and dorsal stream (Leeds et al., 2013). Beside the earliest fMRI habituation studies reporting distance-dependent signal release from adaptation in the intraparietal sulcus, later studies using MVPA and pRF found that numerosity adaptation changed the pattern of number-related activity read out from parietal areas (Castaldi et al., 2016) as well as the preferred numerosity within several numerosity maps located in the temporo-occipital junction, in the superior end of the parieto-occipital sulcus, in the postcentral sulcus and at the junction of the precentral and superior frontal sulci (Tsouli et al., 2021).

Using a habituation paradigm, one seminal study by Izard et al. (2009) in human infants found that shape deviants modulated the ERP responses in ventral temporal areas, whereas number deviants modulated the signal in a right parieto-prefrontal network. Importantly however, the authors observed that number deviants modulated the activity in temporal regions as well. Specifically, they found an antagonistic relationship where number deviants decreased activation in the left anterior temporal regions responsive to object deviants and increased activation in the same region in the right hemisphere, suggesting that object shape (or identity) and numerosity might interact at this level. While mounting evidence shows that the numerical information is also represented along the ventral pathway (Cai et al., 2021, 2023; Karami et al., 2023) future studies should further investigate where adaptation to numerosity and object identity interact.

In conclusion, our findings indicate that numerosity perception can be modulated by contextual effects with previous stimuli capable of distorting numerosity estimates of the following one presented in the same location. Part of the magnitude of these adaptation aftereffects turned out to be selective for salient non-numerical low-level features likely providing cues to define objects’ identity. The results also suggest that numerosity adaptation is likely to occur at relatively high processing levels and that it is influenced by other processes relevant to numerosity perception, including object segmentation and grouping.

## ACKNOWLEDGMENTS

This research was funded by the European Union - Next GenerationEU (PRIN 2022, Project ‘RIGHTSTRESS—Tuning arousal for optimal perception’, Grant no. 2022CCPJ3J, CUP: B53D23014530001) and by the European Research Council (ERC) under the European Union’s Horizon 2020 research and innovation programme (grant n. 832813, ERC Advanced “Spatio-temporal mechanisms of generative perception —GenPercept”, CUP: B14I19000840006).

